# Prestimulus neural alpha power predicts confidence in discriminating identical auditory stimuli

**DOI:** 10.1101/285577

**Authors:** Malte Wöstmann, Leonhard Waschke, Jonas Obleser

## Abstract

When deciding upon a sensory stimulus, the power of prestimulus neural alpha oscillations (~10 Hz) has been shown to hold information on a perceiver’s bias, or confidence, as opposed to perceptual sensitivity per se. Here, we test whether this link between prestimulus alpha power and decision confidence previously established in vision and somatosensation also holds in the auditory modality. Moreover, confidence usually depends on the physical evidence available in the stimulus as well as on decision accuracy. It is unclear in how far the link between prestimulus alpha power and confidence holds when physical stimulus evidence is entirely absent, and thus accuracy does not vary. We here analysed electroencephalography (EEG) data from a paradigm where human listeners (N = 17) rated their confidence in the discrimination of the pitch of two tones that were, unbeknownst to the listeners, identical. Lower prestimulus alpha power as recorded at central channel sites was predictive of higher confidence ratings. Furthermore, this link was not mediated by auditory evoked activity. Our results support a direct link between prestimulus alpha power and decision confidence. This effect, first, shows up in the auditory modality similar to vision and somatosensation, and second, is present also in the complete absence of physical evidence in the stimulus and in the absence of varying accuracy. These findings speak to a model wherein low prestimulus alpha power increases neural baseline excitability, which is reflected in enhanced stimulus-evoked neural responses and higher confidence.

---

Human perception close to threshold is subject to ongoing changes in brain activity. A prevalent view holds that lower power of prestimulus alpha oscillations (~10 Hz) enhances neural sensitivity and thereby the precision of neural stimulus representation. Evidence for this view comes from studies showing a negative relation between prestimulus alpha power and the probability to detect visual targets (Hanslmayr et al., 2007; van Dijk et al., 2008; Busch et al., 2009), tactile targets (Weisz et al., 2014), and to correctly respond to lateralized targets in visuo-spatial attention tasks (Thut et al., 2006; Kelly et al., 2009). Alternatively, more recent research suggests that lower prestimulus alpha power does not lead to more precise but rather to overall amplified neural representation, which is supported by the negative relation of prestimulus alpha power and subjective perception (Lange et al., 2013), decision confidence (Samaha et al., 2017), perceptual bias (Limbach and Corballis, 2016; Benwell et al., 2017a; Iemi et al., 2017), perceptual awareness (Benwell et al., 2017b), and the self-rated level of attention (Whitmarsh et al., 2017); a host of metrics that quantify judgements of one’s own cognitive and perceptual states and are thus referred to here as “metacognition” (for review, see Fernandez-Duque et al., 2000).

It is at present unclear whether the relation of prestimulus alpha power and metacognitive measures such as confidence holds across sensory modalities. Especially, it is unclear whether this relationship holds in the auditory modality, where alpha power has been shown to behave differently compared to other modalities. Visual and somatosensory tasks typically induce alpha power modulation in cortex regions processing visual and somatosensory information, respectively (i.e., occipital and somatosensory cortex regions). However, auditory tasks often show alpha power modulation in non-auditory, parieto-occipital cortex regions (Foxe et al., 1998; e.g., Fu et al., 2001; Strauß et al., 2014). Thus, if the goal is to compare alpha power modulation between modalities, it is necessary to carefully take into consideration the topographic distribution of observed alpha power modulation. Furthermore, the specific direction of an alpha power modulation is crucial when comparing modalities: Attention to sound elicits parieto-occipital alpha power increases, whereas attention to vision elicits alpha power decreases in these areas. In the present study, we expected an alpha power modulation specific in topography and direction: If the relation of prestimulus alpha power and confidence holds across sensory modalities and for audition in particular, alpha power in auditory regions should correlate negatively with a listener’s confidence in their auditory perceptual decisions.

Furthermore, the impact of prestimulus alpha power on ensuing decision confidence might depend on the presence of evidence in the stimulus, or varying evidence in the stimulus and accuracy in the experimental task: Both confidence and alpha power covary with task ease and thus with task accuracy. In turn, both task ease and task accuracy benefit from a stimulus providing more evidence in favour of one or the other decision. For instance, simultaneity judgements of two tactile events displaced briefly in time revealed a negative relation of confidence with prestimulus alpha power in correct trials, while the relation was instead positive in incorrect trials (Baumgarten et al., 2016).

Previous studies have used fixed stimuli in a subset of trials or statistical control of potential influences of the availability of evidence in the stimulus (within a trial), or varying evidence and accuracy (across trials). However, it must be noted that only certain predefined types of influences (e.g., linear, quadratic) can be controlled for statistically. Furthermore, the mere presence of varying evidence and accuracy across trials of an experiment might affect a participant’s behaviour in various ways. To examine the precise relationship of prestimulus alpha power and decision confidence, it is thus necessary to keep these potential influences entirely constant. Samaha and colleagues (2017) statistically controlled for trial-by-trial variance in accuracy and found a negative link between prestimulus alpha power and confidence. While Limbach and Corballis (2016) found a relationship of prestimulus alpha power and the false alarm rate in trials containing no stimulus, Benwell and colleagues (2017b) found no relationship of prestimulus alpha power and perceptual awareness ratings in a subset of trials that excluded evidence in the stimulus. As a results of these conflicting results it is at present not clear whether the link between prestimulus alpha power and metacognitive measures is contingent on other factors in favour of one decision, such as the availability of stimulus evidence or trial-by-trial variation in stimulus evidence.

The present study tests whether the relationship of prestimulus alpha power and decision confidence extends to situations where no evidence for a perceptual decision is available throughout (see Fig. 1D).

**Figure 1.**
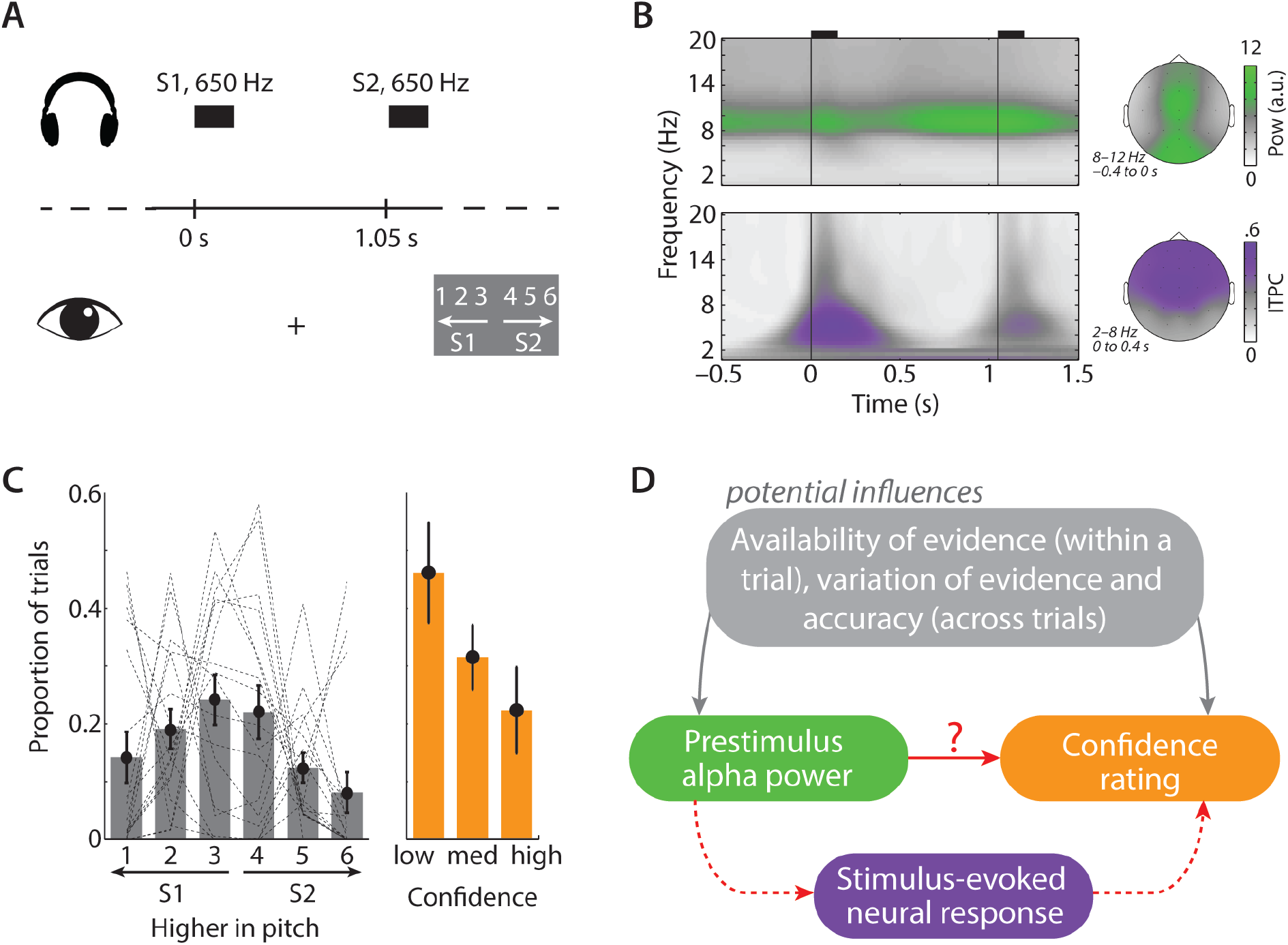
Task design and behavioural and neural variables of interest. (A) Participants had the task to discriminate the pitch of two identical sine tones (650 Hz, 150 ms duration) and to rate their confidence in the decision. A fixation cross was shown throughout the trial. The response screen was only shown in trials including feedback. (B) Grand average absolute oscillatory power (top) and inter-trial phase coherence (bottom) across *n* = 17 participants and 22 scalp electrodes. Topographic maps show the spatial distribution of both measures. (C) Grey bars and dashed lines show average and single-subject proportions of responses, respectively. For further analyses, responses were transformed to low (responses 3 & 4), medium (responses 2 & 5), and high confidence (responses 1 & 6), indicated by orange bars. Error bars show ±1 SEM. (D) While previous research has shown that prestimulus alpha power relates to confidence ratings in the context of varying evidence in the stimulus and varying accuracy (grey box), we asked whether a direct link (red) could be established in a situation where no evidence is present in the stimulus and accuracy does not vary. Furthermore, the present study also tested for a possible indirect link, mediated by the stimulus-evoked neural response (dashed line).

Here, we re-analyse data from a forced-choice pitch discrimination task of two tones (Waschke et al., 2017), which entirely eliminated the presence of evidence and thus variations in evidence and decision accuracy from all trials of the experiment. Unbeknownst to participants, the two tones were physically identical on each trial and thus no evidence in the stimulus in favour of one decision was available. Furthermore, decision accuracy did not vary since participants’ judgments of pitch difference (i.e., first versus second tone higher in pitch) were objectively incorrect throughout, although participants subjectively perceived pitch differences. Importantly, subjective ratings of decision confidence hence were entirely detached from physical evidence and rather driven by fluctuations in brain activity. With these data, we first provide evidence that confidence in auditory decisions relates negatively to prestimulus alpha power as it has been found previously for vision and somatosensation. Second, this relationship holds also in the absence of evidence for decisions. Third, we show that the link from alpha power to decision confidence is a direct one and is not mediated by the intermittent auditory-evoked neural response.

## Materials and Methods

In the present study, we re-analysed data from a previously published experiment (Waschke et al., 2017). Below, we describe essential methodological aspects but refer to the original publication for further details.

### Participants

Data of 17, healthy participants (19–69 years; *M_age_* = 42.65 years; 12 females) were included in the analyses. Data of two additional participants were discarded because they exclusively used the most extreme possible ratings of confidence (i.e., ratings 1 or 6) in pitch discrimination throughout all trials. Participants were financially compensated for participation. The local ethics committee of the University of Lübeck approved all experimental procedures.

### Stimulus materials and task

On each trial of the main experiment, the same sine tone (650 Hz, 150 ms duration, rise and fall times of 10 ms) was presented twice, with an inter-stimulus-interval of 900 ms. Immediately after the offset of the second tone, a response screen was shown (Fig. 1A) until participants entered a response (time limit of 2 s). Participants performed a 2AFC pitch discrimination task with confidence rating. They indicated on each trial which one of two tones was higher in pitch and how confident they were in this decision.

In detail, participants pressed one of 6 buttons, ranging from 1 (first tone clearly higher as second) to 6 (second tone clearly higher as first). Thus, ratings of 1 and 6 corresponded to high confidence, ratings of 2 and 5 to medium confidence, and ratings of 3 and 4 to low confidence. The mapping of response buttons was reversed for 8 of the 17 participants. After an average inter-trial-interval of 3 s (randomly jittered between 2 and 4 s), the next trial started, indicated by the fixation cross changing its colour from grey to light green and back to grey over a period of 500 ms.

Each participant performed 500 trials (except for one participant, who performed 600 trials), divided in blocks of 100 trials each. Bogus feedback was provided for the first few trials of each block (first 10 trials for two participants and first 20 trials for all other participants), where, in 65% of all feedback trials, positive feedback indicating correct pitch discrimination was given. This proportion of positive bogus-feedback was chosen to keep participants engaged in the task. In trials involving bogus feedback, the response screen was followed by a sound indicating a correct or incorrect response after 100 ms. Additionally, after every 20th trial, sham accuracy scores, indicating sham-average accuracy in the past 20 trials, randomly chosen from a uniform [55;65]-% distribution were displayed on the screen for 3 s. For further analyses, trials followed by feedback were excluded.

Before the main experiment, each participant performed 20 practice trials and an adaptive tracking procedure. This procedure was identical to the main experiment but we presented two tones of different pitch on each trial. During the course of the adaptive tracking, the pitch difference was gradually decreased. This was to ensure that participants were in the belief that the two tones in the main experiment were different in pitch, although difficult to discriminate.

### EEG recording and preprocessing

The electroencephalogram (EEG) was recorded at 24 passive scalp electrodes (SMARTING, mBrainTrain, Belgrade, Serbia) at a sampling rate of 500 Hz (DC to 250 Hz bandwidth), referenced against electrode FCz. Electrode impedances were kept below 10 kΩ. The amplifier was attached to the EEG cap (Easycap, Herrsching, Germany) and the EEG data were transmitted via Bluetooth to a nearby computer, which recorded the data using the Labrecorder software (part of Lab Streaming Layer, LSL; Kothe, 2014).

Offline, the continuous data were bandpass-filtered (0.5–100 Hz), re-referenced to the average of both mastoids (which were discarded from all further analyses), and epoched from –2 to +2 s relative to the onset of the first tone (S1). An independent component analysis was used to remove artefact-related components. Remaining artefactual epochs were removed afterwards by visual inspection. All data analyses were carried out in Matlab (R2013b and R2018a), using custom scripts and the Fieldtrip toolbox (Oostenveld et al., 2011).

### Analysis of neural oscillatory signatures

To obtain time-frequency representations of single-trial EEG data, we calculated complex Fourier coefficients for a moving time window (frequency-adaptive Hann-tapers with a width of 4 cycles; moving in steps of 0.01 s through the trial) for frequencies 1– 40 Hz. Oscillatory power was obtained by squaring the magnitude of the Fourier representation. To obtain Inter-trial phase coherence (ITPC), Fourier representations were divided by their magnitudes and averaged across trials, followed by calculating the magnitude of the resulting complex value.

In the present study, we assessed the stimulus-evoked neural response by means of low-frequency ITPC (2–8 Hz) in the first 400 ms following sound onset. ITPC neglects magnitude and polarity of the EEG time domain signal, which might affect other measures of evoked responses, such as the event-related potential (ERP) or evoked power. Thus, an advantage of ITPC is that it can aggregate across evoked response components within the first several hundreds of milliseconds after stimulus onset, which would separate into several more short-lived ERP components of different polarities. Since we did not hypothesize that prestimulus alpha power would affect a particular early stimulus-evoked ERP component but rather evoked activity within the first several hundreds of milliseconds following sound onset, we used ITPC as a measure of the evoked response.

To obtain a single-trial measure of ITPC, we used a Jackknife approach proposed by Richter, Thompson, Bosman and Fries (2015). In brief, conventional ITPC can be obtained for a group of *N* trials but is not defined for a single trial. In order to nevertheless obtain a single-trial metric of ITPC for each participant, we calculated ITPC for all leave-one-out subsamples of trials, resulting in *N* jackknife-ITPC (jITPC) values. If a single trial is highly phase-coherent with remaining trials, leaving this trial out results in a relatively small value of jITPC. Thus, for better interpretability of results, we refer to 1–jITPC as *single-trial phase coherence*. 1–jITPC is a robust measure since it is calculated on a large number of (*N*–1) trials. Nevertheless, differences in 1–jITPC values across trials reliably reflect the relative single-trial phase-locked neural response.

### Statistical analyses

For repeated-measures analyses, we report Greenhouse-Geisser (GG) epsilon (ε) and GG-corrected *p*-values in case of violation of sphericity (*p* < .05 in Mauchly’s test).

The relationship between prestimulus alpha power and auditory-evoked phase coherence on the one hand and decision confidence on the other hand was analysed in two ways. First, for confirmatory analyses, for each participant, single-trial prestimulus alpha power (8–12 Hz; –0.4 to 0 s; 10 central electrodes shown in Fig. 2A), decision confidence (coded as 1, 2, 3 for low, medium, and high confidence, irrespective of whether the decision was made for S1 or S2 as being higher in pitch), and post-stimulus single-trial phase coherence (1–jITPC; 2–8 Hz; 0 to 0.4 s; 10 central electrodes) were extracted. Ten central electrodes (Cz, Fz, Pz, CPz, C3, C4, F3, F4, P3, P4) were chosen as an ROI to focus on alpha power and evoked phase coherence in auditory regions. For each participant, we binned single-trial confidence ratings and single-trial phase coherence according to the magnitude of prestimulus alpha power into four bins (non-overlapping; same trial number across bins), followed by averaging across trials per bin. We then fitted linear functions to model changes in mean confidence ratings and mean single-trial phase coherence as a function of the increasing alpha power bin number (using the polyfit function in Matlab), and tested linear fit coefficients against zero (using one-sample *t*-tests).

**Figure 2.**
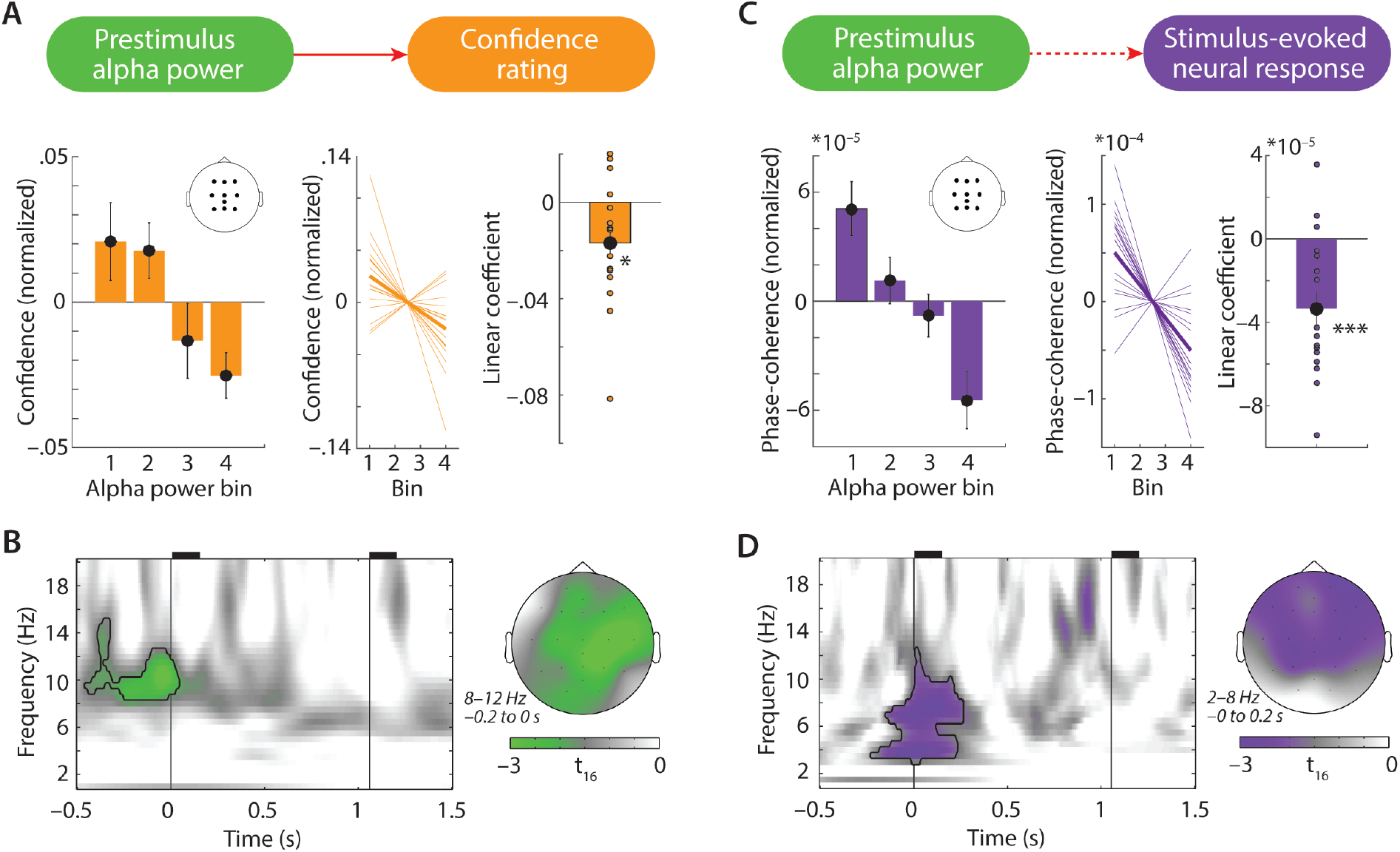
Prestimulus alpha power relates to confidence and auditory-evoked phase coherence. Relationship between prestimulus alpha power and confidence. Left: Bars show mean confidence for four bins of increasing prestimulus alpha power (8–12 Hz; –0.4 to 0 s; 10 central electrodes). Confidence was normalized for each subject by subtraction of average confidence across all alpha power bins. Middle: Orange lines show individual participant’s linear fits to confidence as a function of increasing alpha power bin number. Thick line shows the average fit. Right: Bar and dots shows average and individual linear fit coefficients, respectively, which were significantly smaller than zero; * *p* < 0.05. Error bars show ±1 SEM. (B) Result of a cluster permutation test, which regressed single-trial oscillatory power on single-trial confidence. The black outline indicates a negative cluster (*p* = 0.067), which shows decreasing prestimulus alpha power with increasing confidence. (C) Same as (A) but for stimulus-evoked phase coherence (2–8 Hz; 0 to 0.4 s; 10 central electrodes), which decreased as a function of increasing prestimulus alpha power; * * \**p* < 0.001. (D) A cluster permutation test, which regressed single-trial phase-coherence estimates on single-trial prestimulus alpha power, revealed a significant negative cluster (black outline; *p* < 0.001).

Second, since binning one variable according to the magnitude of a second variable might depend on the number of bins used (Wainer et al., 2006), we backed up our statistical analyses by exploratory analyses (i.e., exploring the entire time-frequency-electrode space for significant effects) using cluster-based permutation test (Maris and Oostenveld, 2007) with continuous (non-binned) predictors. In detail, we applied a two-level statistical approach (Obleser et al., 2012): On the first (single-subject) level, we used an independent samples regression *t*-test, to regress time-frequency representations (–0.5 to 1.5 s relative to S1 onset; 1–20 Hz; all 22 electrodes) of oscillatory power and single-trial phase coherence (1–jITPC) on confidence ratings and single-trial prestimulus alpha power, respectively. This procedure resulted in two time-frequency-electrode spaces of *t*-values for each participant, which were tested against zero using two cluster-based permutation dependent-samples *t*-test on the second (group) level. These tests clustered *t*-values of adjacent bins with *p-*values < 0.05 (minimum cluster size: 2 adjacent electrodes) and compared the summed *t*-statistic of the observed cluster against 10,000 randomly drawn clusters from the same data with permuted condition labels. The *p*-value of a cluster corresponds to the proportion of Monte Carlo iterations in which the summed *t*-statistic of the observed cluster is exceeded (one-tailed).

### Bayes Factor analysis and Effect sizes

For ANOVAs we calculate the *Bayes Factor* (*BF*), using R studio (Version 1.0.136) and the *BayesFactor* package with the default parameters implemented in the *anovaBF* function. For *t*-tests and Pearson correlations we calculate the BF using JASP (version 0.8.1.1). In essence, a *BF* close to 1 indicates that the data are equally plausible under the null and alternative model, while *BFs* < 0.33 begin to lend support to the null model, and *BFs* > 3 begin to lend support to the alternative model (Jeffreys, 1939/1961).

As effect sizes, we report partial eta-squared (*η^2^_p_*) for repeated-measures ANOVAs, and *r-*equivalent (bound between 0 and 1; Rosenthal and Rubin, 2003) for *t*-tests.

## Results

### Neural oscillatory dynamics and confidence in the discrimination of identical tones

Before investigating the relationship between neural oscillatory dynamics and participants’ confidence ratings, we performed descriptive analyses on both of these measures, separately. The neural measures of interest in this study were prestimulus alpha power (8–12 Hz, –0.4 to 0 s) and auditory-evoked low-frequency phase coherence (2–8 Hz, 0 to 0.4 s), which were most prominent at respective centro-parietal and fronto-central electrodes (Fig. 1B).

At the end of each trial, participants judged which one of two tones they perceived as being higher in pitch and how confident they were in this decision, by pressing one button on a 6-point scale, ranging from 1 (first tone, S1, clearly higher in pitch) to 6 (second tone, S2, clearly higher in pitch). Average proportions of responses did not differ significantly for the six response options (Fig. 1C; repeated-measures ANOVA; Greenhouse-Geisser *ε* = 0.39; *F_5,80_* = 2.08; *p* = 0.142; *η^2^_p_* = 0.12; *BF* = 1.36).

As we reported before in Waschke et al. (2017), responses 1–3 (indicating higher pitch of S1) were relatively more frequent than responses 4–6 (indicating higher pitch of S2; *t_16_* = 2.89; *p* = 0.01; *r* = 0.59; *BF* = 5.1). This indicates a general bias to judge the first of two identical tones as being higher in pitch.

For the purpose of the present analyses, responses were converted to low confidence (responses 3 & 4), medium confidence (responses 2 & 5), and high confidence (responses 1 & 6), irrespective of whether S1or S2 was perceived as being higher in pitch (Fig. 1C, orange bars).

### Prestimulus alpha power predicts confidence in pitch discrimination

The major objective of this study was to test for a direct link of prestimulus alpha power and confidence in auditory decisions in a task without potential influences of varying evidence in the stimulus or varying accuracy (red solid line in Fig. 1D). Indeed, our results support the existence of such a link (Fig. 2A): With increasing levels of prestimulus alpha power (8–12 Hz; –0.4 to 0 s; 10 central electrodes) confidence decreased (*t_16_* = –2.73; *p* = 0.015; *r* = 0.56; *BF* = 3.91; significant also for 3 and 5 bins: *ps* < 0.025; *rs* > 0.5; *BFs* > 3; linear change in confidence across 4 bins of alpha power not correlated with participants’ age, *r* = 0.12; *p* = 0.646; *BF* = 0.33). Note that this effect was also present, albeit weaker, when alpha power was instead obtained from a single-window spectral estimate calculated exclusively from prestimulus (– 0.4 to 0 s) EEG time-domain data (*t_16_* = –1.98; *p* = 0.065; *r* = 0.44; *BF* = 1.2), which rules out the possibility that this effect is driven by post-stimulus EEG activity.

Next, we controlled for potential influences of non-normality of single-trial alpha power values and linear change of alpha power (and confidence) across the duration of the experiment (Benwell et al., 2018). To this end, we first log-transformed single-trial alpha power values and, second, removed the linear change in alpha power and confidence across trial number, using the residuals of two separate linear regressions of alpha power and confidence on trial number. The negative relation of residuals of prestimulus alpha power and residuals of decision confidence remained significant (*t_16_* = –2.81; *p* = 0.013; *r* = 0.58; *BF* = 4.48; significant also for 3 and 5 bins: *ps* < 0.025; *rs* > 0.5; *BFs* > 2.7).

To furthermore explore the specificity of the relation of alpha power and confidence in time-frequency-electrode space, we performed a cluster permutation test to regress single-trial power on single-trial confidence ratings. The test revealed one negative cluster close to statistical significance (Fig. 2B; cluster *p*-value = 0.067), which was limited to the prestimulus time range and to the alpha frequency band. No additional significant clusters were found (all *ps* > 0.25). Critically, no significant cluster was found prior to second tone (S2). This suggests that in the present paradigm with a relatively short inter-stimulus-interval between the two tones, the relation of neural alpha power and confidence only holds for the time interval prior to the onset of the first one of two tones.

Despite the fact that the extent of a cluster in time-frequency-electrode space depends on various parameters of the cluster test and should be interpreted with care (Maris and Oostenveld, 2007), the temporal extent of the significant prestimulus cluster in Fig. 2B clearly suggests that it does not result from temporal smearing of post-stimulus activity due to the width of the time window used for time-frequency analysis: First, the significant cluster includes virtually only prestimulus time points, whereas temporal smearing should be symmetrical and would thus smear a post-stimulus effect also into the post-stimulus time range. Second, the cluster starts more than 200 milliseconds before stimulus onset and thus before the earliest time point that might be affected by temporal back-smearing of post-stimulus activity. (The analysis window with a width of 4 cycles centred at stimulus onset ranges from –200 to +200 ms for a 10-Hz alpha oscillation.)

### The influence of prestimulus alpha on confidence is not mediated by stimulus-evoked activity

Note that the most important result of the present study, that is, the negative relation of prestimulus alpha power and confidence, could be an indirect one. The effect of prestimulus alpha power on confidence might be entirely or partly mediated by the intermittent stimulus-evoked neural response (dashed line in Fig. 1D) that follows upon such a prestimulus alpha state.

A first, necessary but not sufficient, precondition for such a mediation (Baron and Kenny, 1986) would be a significant relation between prestimulus alpha power (8–12 Hz; –0.4 to 0 s; 10 central electrodes) and the stimulus-evoked response (2–8 Hz; 0 to 0.4 s; 10 central electrodes). This was the case. We binned the stimulus-evoked response for alpha power, which revealed a significant negative relationship (Fig. 2C; *t_16_* = –4.11; *p* < 0.001; *r* = 0.72; *BF* = 44.93; significant also for 3 and 5 bins: *ps* < 0.005; *rs* > 0.6; *BFs* > 10). Again, we controlled for non-normality of single-trial alpha power and linear changes of alpha power (and the stimulus-evoked response) across the duration of the experiment. To this end, we first log-transformed single-trial alpha power values and, second, removed the linear change in alpha power and single-trial phase coherence across trial number, using the residuals of two separate linear regressions of alpha power and single-trial phase coherence on trial number. The negative relation of residuals of prestimulus alpha power and residuals of the stimulus-evoked neural response remained significant (*t_16_* = –4.01; *p* = 0.001; *r* = 0.71; *BF* = 37.55; significant also for 3 and 5 bins: *ps* ≤ 0.002; *rs* > 0.66; *BFs* > 20). Furthermore, the negative relation of alpha power and the stimulus-evoked neural response remained significant when alpha power was estimated from a single-window spectral estimate calculated on only prestimulus data (– 0.4 to 0s) and the stimulus-evoked neural response was estimated from a single-window spectral estimate calculated on only post-stimulus data (0 to 0.4s; *t_16_* = –4.14; *p* < 0.001; *r* = 0.72; *BF* = 47.74).

To further explore the specificity of the relation of prestimulus alpha power and the evoked response in time-frequency-electrode space, we regressed single-trial phase coherence on single-trial prestimulus alpha power. A cluster permutation test confirmed the negative effect of prestimulus alpha power on the stimulus-evoked phase coherence in low frequencies (Fig. 2D; significant cluster *p*-value < 0.001). No additional significant clusters were found (all *ps* > 0.4).

A second necessary precondition for the stimulus-evoked response as a mediator of the prestimulus alpha-confidence relation would be a substantial reduction of this relation under statistical control for the stimulus-evoked response (e.g., Baron and Kenny, 1986).

This, however, was not the case, ruling out a mediated relationship. In detail, we first eliminated variance in prestimulus alpha power and confidence explained by the stimulus-evoked response through regression of these two variables on single-trial phase coherence, using two linear regressions for each participant. We then performed the same binning analysis used before to model residuals of confidence ratings for the binned residuals of alpha power, which again yielded a significant negative relation (*t_16_* = –2.78; *p* = 0.013; *r* = 0.57; *BF* = 4.25; significant also for 3 and 5 bins: *ps* < 0.015; *rs* > 0.55; *BFs* > 4.8). Thus, the relationship between prestimulus alpha power and confidence was not reduced under control for the stimulus-evoked response, and thus not mediated by it. Using the same procedure, we controlled for a possible mediation of the alpha power-confidence relationship by the difference in the evoked response to S1 minus S2, which was not the case (i.e., the alpha power-confidence relationship was still significant when we regressed out the influence of the difference in the evoked response to S1 minus S2; *t_16_* = –2.93; *p* = 0.01; *r* = 0.59; *BF* = 5.46).

We also tested whether the stimulus-evoked response was related directly to confidence. Binning of confidence for single-trial stimulus-evoked phase coherence revealed no significant linear relationship (*t_16_* = –0.97; *p* = 0.347; *r* = 0.24; *BF* = 0.38; non-significant also for for 3 and 5 bins; *p*s > 0.2; *r*s < 0.35, *BFs* < 0.55). Neither did a cluster permutation test regressing single-trial phase coherence on single-trial confidence (all cluster *p*-values > 0.15).

### Prestimulus alpha power does not predict decision outcome

Finally, it might be that prestimulus alpha power is not only related to confidence but also to the actual pitch discrimination outcome (i.e., experience of the first vs. the second tone as being higher in pitch). We performed two analyses to test this. First, binning the proportion of decisions in favour of S1 as being higher in pitch according to prestimulus alpha power revealed no significant linear relationship (*t_16_* = –0.92; *p* = 0.37; *r* = 0.22; *BF* = 0.36; non-significant also for 3 and 5 bins: *ps* > 0.3; *rs* < 0.3; *BFs* < 0.4). This relationship remained non-significant when repeating the analysis with log-transformed alpha power values and removal of the linear change in alpha power and the decision outcome across trial number (*t_16_* = – 0.94; *p* = 0.362; *r* = 0.23; *BF* = 0.37; non-significant also for 3 and 5 bins: *ps* > 0.3; *rs* < 0.27; *BFs* < 0.39).

Second, a cluster permutation test to regress oscillatory power on decisions for S1 versus S2 as being higher in pitch did not reveal any significant clusters (all *p*s > 0.3).

## Discussion

Recently, evidence has accumulated that prestimulus alpha power might influence metacognitive measures in the aftermath of a stimulus, such as confidence in perceptual decisions close to threshold. Here, we first demonstrate that this effect, and notably its direction, surfaces in the auditory modality just as it does for vision and somatosensation. Second, this relation does not hinge on potential influences of the availability of evidence in the stimulus or varying evidence and accuracy across trials but persists in the most extreme cases of perception, that is, in the entire absence of physical evidence. Third, this relation is a direct effect of alpha power on confidence, as it is not simply mediated by differences in the stimulus-evoked neural response. These findings lend plausibility and parsimony to the suggested mechanistic role of alpha oscillations in regulating neural baseline excitability.

### Prestimulus alpha power links to confidence in auditory decisions

Prior studies to demonstrate a relationship between prestimulus alpha power and confidence used visual (Samaha et al., 2017) or somatosensory (Craddock et al., 2017) tasks (for evidence of prestimulus influences on auditory perception, see Kayser et al., 2016). In one previous study in the auditory modality, we found that participants’ confidence in speech comprehension was negatively related to alpha power. This however occurred post- not prestimulus onset and in a task where confidence and accuracy did covary strongly (Wöstmann et al., 2015).

It might seem unsurprising that the present study conceptually replicates in the auditory modality previously shown negative links of prestimulus alpha power and decision confidence in vision and somatosensation. However, the net alpha power measured in human scalp EEG is clearly dominated by visual, that is occipito-parietal alpha. This is reflected in pervasive maximal alpha power modulation in occipito-parietal regions, even in auditory attention and memory tasks (e.g., Lim et al., 2015; Wöstmann et al., 2015, 2017). Compared to tasks in the visual modality, auditory tasks often reverse the modulation of visual alpha power rather than exhibiting an effect on auditory alpha power (e.g., Fu et al., 2001; Strauß et al., 2014).

Furthermore, existence and function of spontaneous alpha oscillations in auditory regions is a matter of debate (Lehtelä et al., 1997), although evidence in favour of auditory alpha generators has been demonstrated by source-projected spectral activation profiles (Keitel and Gross, 2016), human electrocorticographic recordings from auditory cortical regions (Gomez-Ramirez et al., 2011; for a review see e.g. Weisz et al., 2011) and neuroelectric recordings in monkeys (Lakatos et al., 2016). In the present study, maximum prestimulus alpha modulation in relation to confidence was observed at central electrodes (Fig. 2B). In a post-hoc analysis of the present results (not shown), we found that the negative links between prestimulus alpha power and confidence as well as the stimulus-evoked response were not significant when these neural responses were extracted at occipital (O1, O2, POz) instead of central electrodes. This is at least circumstantial evidence against a purely visual or supramodal parietal alpha power modulation (Banerjee et al., 2011) and rather speaks to alpha power modulation in sensory-specific, auditory regions.

### Direct link between prestimulus alpha power and confidence

Although our participants heard two instances of the very same tone on each trial, many of them reported perception of pronounced pitch differences when debriefed after the experiment and did not raise concerns regarding the true nature of our stimuli. Together with previous work on perception of differences between identical stimuli, this speaks to the feasibility of such a task structure (Amitay et al., 2006, 2013; Bernasconi et al., 2011). Note that it is controversial what factors make a participant report high (versus low) confidence in a decision: Confidence likely reflects a participant’s subjective experience that the made decision is correct, given the evidence (Pouget et al., 2016). However, confidence has also been found to depend largely on the information in support of the choice made, while information in support of the alternative choice option is largely disregarded (Peters et al., 2017).

The most important finding of the present study is the substantial negative relation of a participant’s prestimulus alpha power on the one hand and confidence in the pitch discrimination of two identical tones on the other (Fig. 2A&B). A similar prestimulus alpha power-confidence relationship has been established before only in the context of available evidence in the stimulus, or varying evidence in the stimulus and varying task accuracy across trials, which, in turn, typically covary with confidence. While previous studies have aimed for statistical control of these potential influences, we eliminated these altogether by reducing the physical evidence for both response alternatives to zero at all times. Our results substantiate that prestimulus alpha power relates directly to decision confidence in the absence of evidence, variations of evidence, or varying accuracy and thus emphasize a more general relation of prestimulus alpha-power and meta-cognitive processes.

Although no statistical inference about the direction of the link between prestimulus alpha power and confidence can be made based on our correlational results, the natural order of these events within a trial speaks to an effect of prestimulus alpha power on confidence. Future studies might investigate the direction of this link more directly, for example by testing the effect of transcranially modulated alpha oscillations (using transcranial alternating current stimulation; Herrmann et al., 2013) on decision confidence. As a note of caution, however, a recent study found an experimental modulation of response criterion to result in modulation of prestimulus alpha power (Kloosterman et al., 2018), which might suggest that the observed link is in fact bi-directional.

Our results somewhat diverge from a recent study by Benwell et al., (2017b). There, the negative relation between prestimulus alpha power and perceptual awareness ratings in a luminance discrimination task decreased with smaller degrees of evidence in the stimulus, and it even disappeared for trials in which no stimulus (and thus no evidence) was presented. The present study exclusively contained trials without evidence available in the stimulus (except for the pre-experiment adaptive tracking procedure). Thus, our participants likely adapted to this situation such that even small subjectively experienced pitch differences (Micheyl et al., 2009) were sufficient to induce considerably varying levels of confidence. Such an adaptation is arguably less likely if trials with relatively large degrees of evidence in the stimulus are included in the experiment.

Of note, the negative relationship of alpha power and decision confidence was found only in the time interval preceding the first but not the second tone (Fig. 2B). In theory, it might be that in experimental paradigms with longer inter-stimulus-intervals (e.g., Iemi and Busch, 2018) alpha power prior to the first versus prior the second stimulus relates to metacognitive measures in sensory decisions. In the present paradigm, however, the two tones were displaced in time by only 900 ms, which likely resulted in generally reduced dynamics of alpha power prior to the second versus the first tone (for a similar argument, see Waschke et al., 2017). Thus, by design, the possibility of observing significant power modulations by confidence prior to the second tone might have been lowered.

### A mechanistic role for prestimulus alpha power in perception

In line with prior research (e.g., Brandt and Jansen, 1991; Becker et al., 2008) we found a negative relation between prestimulus alpha power and the stimulus-evoked neural response, assessed here by single-trial phase coherence (Richter et al., 2015). This finding agrees with the proposed inhibitory role of high alpha power (Jensen and Mazaheri, 2010; Foxe and Snyder, 2011; Strauß et al., 2014), which is thought to decrease neural excitability, reflected in a reduced response to sensory stimulation. It has been proposed that the neural sensory response scales quadratically (i.e., inverted U-shape) with signatures of neural sensitivity, such as prestimulus alpha power (Rajagovindan and Ding, 2011; Kloosterman et al., 2018). The linear relation observed here (Fig. 2C) does not speak against such a quadratic relationship per se; it might be that the range of prestimulus alpha power values was too small to reveal the full quadratic effect.

According to an adapted signal-detection model (Iemi et al., 2017; Samaha et al., 2017) higher neural baseline excitability for sensory discrimination does not increase the difference in the neural representation of to-be-discriminated stimuli, but rather increases the overall neural representation and thus evidence for both decision outcomes.

Behavioural work has demonstrated how stimulus intensity relates to confidence: In two experiments, Zylberberg et al., (2012) revealed that “confidence was influenced by evidence for the selected choice but was virtually blind to evidence for the non-selected choice”^1^. In the present study, lower prestimulus alpha power likely enhanced neural baseline excitability, which increased the evidence for both decision outcomes (see Fig. 3). This enhanced evidence subsequently increased the confidence for the selected choice (i.e., first versus second tone being higher in pitch).

**Figure 3.**
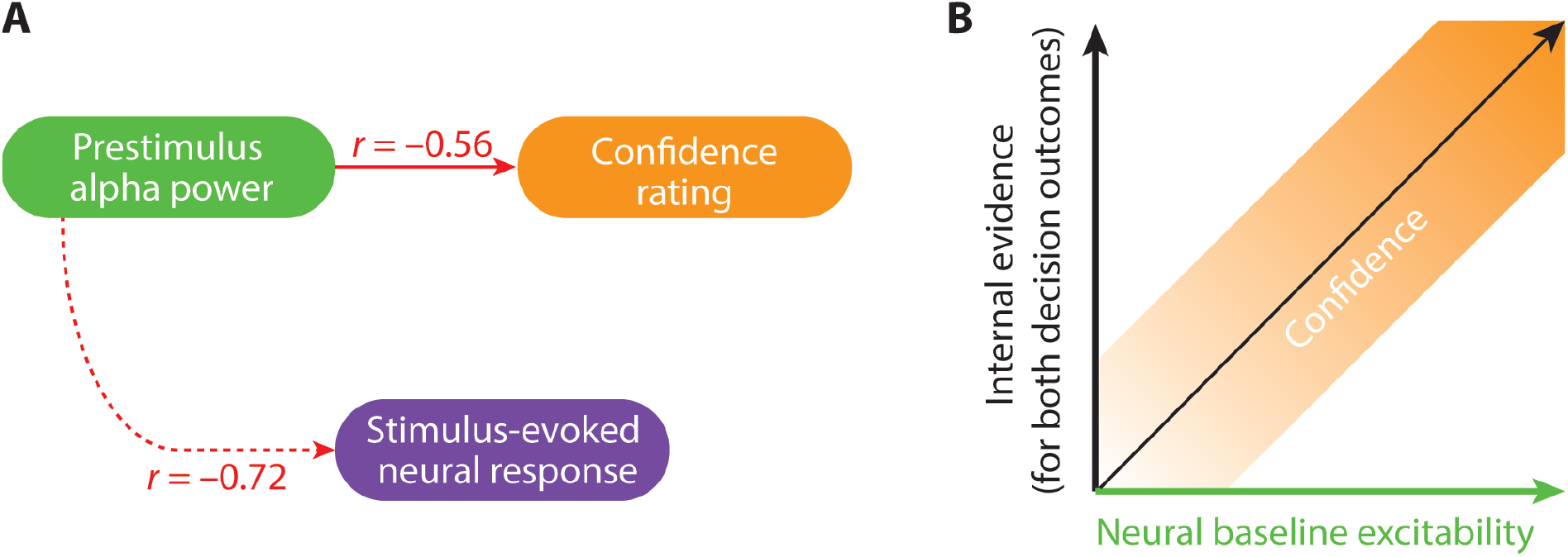
Results of present study and underlying mechanistic relations. (A) Graphical summary of the obtained results. Prestimulus alpha power significantly relates to confidence and to the stimulus-evoked response. Negative signs of *r*-equivalent effect sizes are to indicate that both of these relations were negative. Note that the link between the stimulus-evoked neural response and confidence is omitted as it was weak and not significant (*r* = 0.24; *p* = 0.347). Arrows do not imply directed (or causal) influences but rather temporal succession. (B) A simple mechanistic model to explain the observed results. Prestimulus alpha power is thought to reflect the inverse of neural baseline excitability. With increasing baseline excitability (i.e., decreasing prestimulus alpha power), the overall neural representation of the stimulus is amplified, which increases the participant’s internal evidence for both decision outcomes (i.e., decision of the first versus second tone as being higher in pitch). Since the confidence in a perceptual decision scales positively with the evidence for the selected choice, confidence increases with higher degrees of evidence.

Of note, our results disagree with the alternative view, namely that prestimulus alpha power would relate to the precision of neural representations. According to this view, low prestimulus alpha power should lead to more precise neural representation of the two identical tones. This, in turn, should surface in lower confidence when participants are forced to make a choice regarding the pitch difference, which is the opposite of what we observed here.

Although higher baseline excitability surfaces in increased measures of metacognition such as confidence in several paradigms including the present one, effects on task accuracy or perceptual sensitivity are still possible. For instance, during spatial attention, two sources of alpha power can be differentiated in the contra-versus ipsilateral hemisphere, which are thought to modulate baseline excitability for respective stimuli on the attended versus ignored side (e.g., Worden et al., 2000; Haegens et al., 2011; Wöstmann et al., 2016). With such separate alpha sources, low contra- and high ipsilateral alpha power enhance excitability for the attended as opposed to the ignored stimulus, which increases evidence for the attended stimulus only (eventually resulting in more accurate stimulus selection). A recent study supports this view (Wöstmann et al., 2018): Transcranial stimulation of alpha (versus gamma) oscillations in the left hemisphere decreased recall accuracy of auditory targets presented on the right side.

## Conclusions

To assess the mechanistic relevance of patterns of neural activity in general and of alpha oscillations in particular, we must relate these as closely as possible to changes in human behaviour. We here demonstrate that prestimulus alpha power directly predicts auditory decision confidence, and that this link does not depend on changing evidence in the physical stimulus or on changes in accuracy. These results support a model of cortical alpha oscillations as a proxy for neural baseline excitability that holds across sensory modalities, including audition. In this model, prestimulus alpha power does not lead to more precise but rather to overall amplified neural representations.

## Acknowledgements

Research was supported by the Volkswagen foundation (to JO; BIT-CHAT) and the European Research Council (to JO; ERC-CoG-2014-646696).

## Data accessibility

All data are available from the corresponding authors upon request.

## Footnotes

1 Cited from Zylberberg A, Barttfeld P, Sigman M (2012) The construction of confidence in a perceptual decision. Front Integr Neurosci 6 Available at: http://journal.frontiersin.org/article/10.3389/fnint.2012.00079/abstract.

